# Transposable element-derived siRNAs control viral disease in Arabidopsis

**DOI:** 10.64898/2026.02.06.704401

**Authors:** L Elvira-González, C Himber, C Matteoli, L Felgines, V Fiess, PM Kienzle, J Hennequart, V Schurdi-Levraud, T Blevins, M Heinlein

**Affiliations:** Institut de biologie moléculaire des plantes (IBMP), CNRS, Université de Strasbourg, France; Université de Bordeaux, INRAE, Bordeaux, France

## Abstract

Crop diseases caused by viruses can result in significant economic losses. However, the mechanisms that lead to disease are not well understood. Meanwhile, metagenomic surveys have revealed that most plants in nature are tolerant to viruses, thus maintain their fitness despite of recurrent infection. Using plants that transition from a diseased state to a tolerant state during virus infection, we demonstrate that tolerance depends on the virus-induced production of 21-nucleotide small interfering RNAs (siRNAs) from transposable elements (TEs). These findings indicate that siRNAs derived from TE sequences play a pivotal role in mitigating the detrimental effects of viral infection on plant health.

## Introduction

Plant viruses are ubiquitous, abundant and ecologically important in the environment (Breitbart & Rohwer, 2005). However, they cause a myriad of challenges for agriculture and food security by causing diseases in cultivated plants leading to global crop losses (Nicaise, 2014). By contrast, wild plants are often ‘tolerant’, that is, free of disease symptoms despite recurrent infection (Roossinck, 2012). Tolerance is a defense strategy that maintains plant fitness while allowing the virus to propagate (Pagan & Garcia-Arenal, 2020). This is in contrast to resistance, which sustains plant fitness by defense mechanisms that reduce viral accumulation, including RNA silencing (Lopez-Gomollon & Baulcombe, 2022). Tolerance is more durable in sustaining host fitness than resistance as it does not cause selective pressure on viruses to escape from plant defenses (Pagan & Garcia-Arenal, 2018, 2020). Tolerance therefore has great potential for controlling plant disease and is also highly relevant for understanding plant-pathogen interactions in wild populations (Pagan & Garcia-Arenal, 2020). However, the molecular determinants of plant tolerance to viruses remain poorly understood.

The virome of plants is dominated by RNA viruses (Dolja *et al*, 2020) such as tobacco mosaic virus (TMV). Certain natural accessions of *Arabidopsis thaliana*, such as ecotype Col-0, are tolerant to TMV and show no or only benign symptoms upon infection, whereas other ecotypes develop disease (Dardick *et al*, 2000). Interestingly, Col-0 plants infected with the TMV-related oilseed rape mosaic virus (ORMV) undergo a change from severe disease to tolerance during development: this disease recovery phenotype is scored by observing the transition from symptomatic growth to asymptomatic growth (Kørner *et al*, 2018). Unlike disease recovery due to resistance mediated by antiviral RNA silencing (Baulcombe, 2004; Paudel & Sanfacon, 2018), disease recovery in ORMV-infected plants is not associated with significant changes in viral RNA or viral protein accumulation (Kørner *et al*., 2018; **Fig. 1**), thus reflecting tolerance. The onset of plant tolerance to ORMV leading to disease recovery occurs in emerging leaves and coincides with a loss of the ability of the viral suppressor of RNA silencing (VSR) to suppress GFP transgene silencing in these leaves (Kørner *et al*., 2018). The underlying mechanism depends on small interfering RNA (siRNA) pathway components, including the predominantly cytoplasmic proteins RDR6, SGS3, DCL2, DCL4, HEN1, and AGO1 (Kørner *et al*., 2018). More surprisingly, induction of tolerance also depends on RNA polymerase IV (Pol IV) and RDR2 (Kørner *et al*., 2018), which are core components of the nuclear machinery that generates DCL3-dependent 24 nt siRNAs to guide RNA-directed DNA methylation (RdDM) and silence transposable elements (TEs) (Huang *et al*, 2021; Singh *et al*, 2019; Xie *et al*, 2025). While Pol IV-RDR2 mediate plant recovery from ORMV, downstream RdDM steps involving DCL3, RNA polymerase V (Pol V) and AGO4 are not required (Kørner *et al*., 2018). On the contrary, the induction of tolerance to ORMV is strongly enhanced in *dcl3* null mutants (Kørner *et al*., 2018). Finally, disease recovery was shown to be independent of ethylene, salicylic acid (SA), jasmonic acid (JA) and auxin synthesis or signaling (Kørner *et al*., 2018). Taken together, the available evidence suggests that induction of tolerance in ORMV-infected plants relies on both cytoplasmic and nuclear small RNA pathways. However, the mechanism by which these pathways are activated and the endogenous loci targeted for virus-induced siRNA production remain unknown.

**Fig. 1.**
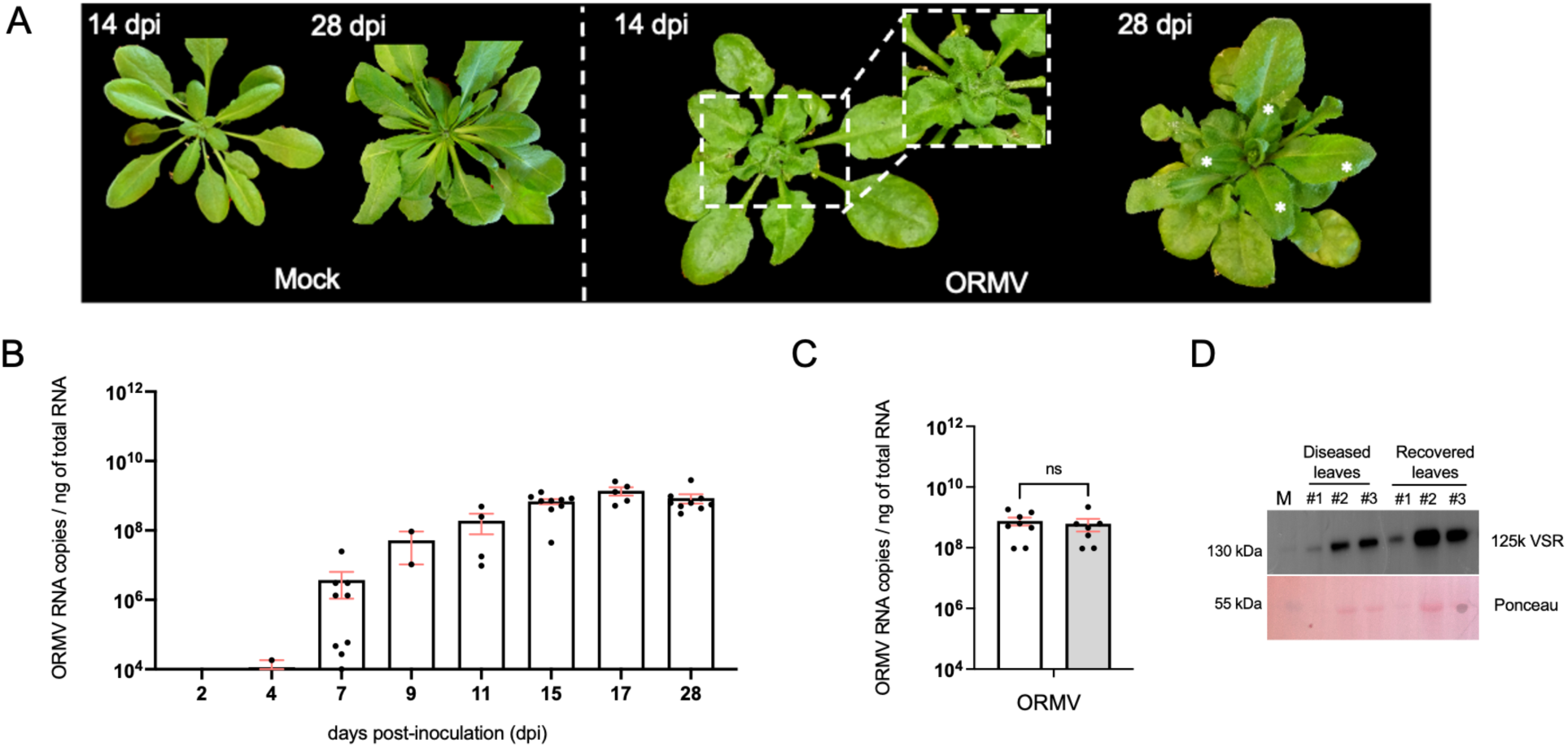
ORMV infection dynamics and disease recovery in Arabidopsis Col-0. (**A**) Mock-inoculated and ORMV-infected plants at 14 and 28 days post-inoculation (dpi). ORMV-infected plants with highly diseased apical leaves at 14 dpi and with diseased and newly developed healthy (recovered) leaves (asterisks) at 28 dpi. (**B** and **C**) ORMV accumulation in infected plants (RT-qPCR). Points represent individual biological replicates. Bars show the mean with standard error (SE). (**B**) Accumulation of ORMV RNA over time in apical leaves. (**C**) The recovered leaves (shaded in grey) accumulate viral RNA to similar levels as the diseased leaves (28 dpi). (**D**) Immunoblot detection of the viral 125-kDa suppressor of RNA silencing (125k-VSR) in diseased and recovered leaves of different plants (#1, #2, and #3) at 28 dpi. Ponceau staining shows the large subunits of Ribulose-1,5-bisphosphate carboxylase/oxygenase (RuBisCo; 51-58 kDa) as a marker for protein loading. M, protein size marker.

## Materials and Methods

### Virus preparation

Virion particles were prepared from ORMV-infected *Nicotiana benthamiana* leaves. Leaves were homogenized to fine powder in liquid N_2_. After addition of 1 mL 0.5 M sodium-phosphate buffer (NaP) pH 7.4 and 0.1% 2-mercaptoethanol per gram leaf material, 1 volume of butanol/chloroform (1/1 (v/v)) was added and the phases separated by centrifugation (twice for 15 min at 12,000 g). Virions in the upper, aqueous phase were precipitated with 4% PEG 8000 at 20,000 g. The pellet was resuspended in 10 mM NaP pH 7.4 and cleared by centrifugation at 5000 g for 10 min. The supernatant was precipitated again with 4% PEG 8000 and 1% NaCl and resuspended in 10 mM NaP, pH 7.4. Virion concentrations were estimated from absorbance values at 260 nm.

### Plant material

*Arabidopsis thaliana* ecotypes Columbia-0 (Col-0, NASC reference N60000) and Shadara (Sha, NASC reference N29735) were used for all experiments. To obtain *CLSY* and *NRPD1* deletion mutants via CRISPR/Cas9 genome editing, single guide RNAs (sgRNAs) were designed using the Geneious Prime “Find CRISPR Sites” functionality with default parameters. sgRNA candidates with high predicted activity scores and maximum specificity in *A. thaliana* were cloned into appropriate shuttle vectors (Stuttmann *et al*, 2021) (Addgene #100000071). For *clsy1234-C9* quadruple mutants, two shuttle vectors containing sgRNA expression cassettes were prepared per *CLSY* gene, then this 8x assembly was cloned into a zCas9io plant transformation vector with plant selection markers (Stuttmann *et al*., 2021). The *nrpd1-1*^Sha^ (*pol IV-C9^Sha^*) mutant was obtained using two sgRNA cassettes cloned as a 2x assembly in the zCas9io plant transformation vector. After validating the final clones by MiSeq whole-plasmid sequencing, each Cas9 editing vector was transfected into *A. tumefaciens* strain GV3101 and used to transform *A. thaliana* ecotypes Col-0 or Sha via the floral dip method. T1 transformants were positively selected by screening fluorescent red seeds. Genomic DNA extraction was isolated for PCR amplification with primers (**Table S5**) flanking putative edit sites. Deletion events were detected by monitoring the occurrence of smaller PCR bands compared to wild-type samples in agarose gel electrophoresis. Sanger sequencing was then conducted using these same primers to precisely determine the length and position of each deletion in the Cas9-generated alleles. Homozygous quadruple *clsy1234-C9*^Col^ and *clsy1234-C9*^Sha^ mutant plants were identified, then backcrossed to the corresponding wild-type Col-0 or Sha parental lines to generate F1 plants and then F2 seeds that segregate *clsy* mutations. Screening for single, double and triple *clsy* mutants was conducted in these F2 populations with a new set of PCR primers (**Table S5**). Other *A. thaliana* mutants used in this study included *clsy1-7 clsy2-2 clsy3-1 clsy4-1* (*clsy1234-TD*^Col^) (Zhou *et al*, 2018), *nrpd1-3* (*pol IV-TD^Col^*) (SALK_128428) (Onodera *et al*, 2005) and *dcl3-1* (SALK_005512) (Xie *et al*, 2004).

### Plant growth conditions and inoculation with virus

Plants were grown in a plant growth chamber at 21°C with 12 h/12 h light/dark cycles. The fifth and sixth leaves of four-week-old plants were each rub-inoculated with 100 ng virions in water or, for mock control inoculations, with water alone. From 15-18 dpi onwards, plants were monitored for the emergence of recovered leaves.

### GUS staining

Plant tissues were harvested and immediately immersed in GUS staining solution ensuring complete submersion of the samples. The GUS staining solution was freshly prepared by mixing equal volumes (1:1) of two solutions: (I) a 2 mM 5-bromo-4-chloro-3-indolyl β-D-glucuronide (X-Gluc) solution containing 50 mM sodium phosphate buffer (pH 7.0), 0.1% (v/v) Triton X-100, and 10 mM EDTA, and (II) a solution containing 1 mM potassium ferrocyanide and 1 mM potassium ferricyanide in 50 mM sodium phosphate buffer (pH 7.0). To facilitate penetration of the staining solution, samples were vacuum infiltrated in an airtight vacuum chamber by gradually reducing the pressure to −0.08 MPa. Following infiltration, plates were sealed, protected from light, and incubated at 37°C for 24 h. The staining reaction was stopped by replacing the staining solution with sodium phosphate buffer, followed by stepwise destaining in increasing concentrations of ethanol (50%, 70%, and 100%, v/v) to remove chlorophyll and preserve tissue structure.

### Viral RNA quantification with TaqMan RT-qPCR

Total RNA was extracted from frozen, ground plant tissue using TRIzol reagent (Invitrogen) according to the manufacturer’s instructions. RNA concentration and purity were determined spectrophotometrically using a NanoDrop instrument (Thermo Scientific). Quantitative reverse transcription PCR (RT-qPCR) was performed using 10 ng/μL of total RNA with the qScript™ XLT One-Step RT-qPCR ToughMix (Quantabio), following the manufacturer’s recommendations. Absolute quantification of viral RNA was achieved using serial dilutions of in vitro-transcribed viral RNA of known copy number, which served as a standard curve to determine the relationship between cycle threshold (Ct) values and transcript abundance. RT-qPCR reactions were carried out under standard cycling conditions, and all reactions included no-template controls to monitor contamination. Viral RNA copy numbers in plant samples were calculated by interpolation from the standard curve. More than three biological replicates were used in each case. Data are presented as mean ± standard error.

### *ONSEN* RT-qPCR

Total RNA was extracted with TRIzol reagent, treated with DNase I (Thermo Scientific) and purified using phenol-chloroform and ethanol precipitation. Reverse transcription was performed using SuperScript IV Reverse Transcriptase (Invitrogen), RiboLock RNase Inhibitor (Thermo Scientific) and random hexamer primers (Thermo Scientific) on 1 μg of RNA previously treated with DNase I. qPCR was done with the LightCycler 480 II (Roche) using Takyon No ROX SYBR 2X (Eurogentec), the cDNA and specific primers to detect *ONSEN* or *UBQ10* transcripts (**Table S5**).

### Northern blot analysis

Total RNA was extracted from ground tissue using TRI Reagent (Sigma-Aldrich). Total RNA was quantified by spectrophotometric analysis using a Nanodrop device (Thermo Scientific). Resulting RNA samples were dehydrated using a SpeedVac (SPD111V, Thermo Scientific), then loaded onto a 16% polyacrylamide gel and run for 1 h at 15 W. The RNA was transferred to Hybond-N+ nylon membrane (GE Healthcare) at 300 mA for 2 h at 4 °C and UV-crosslinked (140 mJ/cm²). The membrane was washed with 50 mL 2X SSC (Saline sodium citrate), prehybridized for 3 h in 20 mL of PerfectHyb Plus hybridization buffer (Sigma-Aldrich), and then hybridized overnight at 35-40 °C with a polynucleotide kinase (PNK) P^32^ end-labeled DNA oligo probe. The membrane was washed three times with 20 mL of 0.5% SDS, 2X SSC and then exposed to a phosphor storage screen for seven days. The signal on the screen was scanned using a Typhoon biomolecular imager (Amersham, GE Healthcare). To rehybridize the membrane, the previous probe was removed by incubating the membrane in hot 0.1% SDS solution (85-95°C) twice for 20 min, and the membrane was then prehybridized in 20 mL of PerfectHyb Plus hybridization buffer before adding the new probe.

### Western blot analysis

Total protein extracts were isolated by grinding plant leaf tissues in liquid N_2_ followed by homogenization in 4x Laemmli buffer (Roche) and heating for 5 min at 95 °C. After centrifugation at 12,000 g for 3 min, samples were added to the gel. Proteins were separated in 12% SDS-PAGE gels, followed by wet blotting to PVDF membranes. The 125k VSR of ORMV was detected with a rabbit antibody raised against a synthetic peptide (GITRADKDNVRTVDS). As comparison for loading, Ponceau staining was used. Signals were detected by luminescence labelling of the primary antibodies with HRP-conjugated secondary antibodies (Thermo Fisher Scientific).

### Cell fractionation

Nuclei were isolated from non-infected, ORMV-infected, and recovered leaves. Leaf tissues were frozen in liquid nitrogen and ground to a fine powder. The powder was resuspended in ice-cold LVM1 buffer (0.4 M sucrose, 10 mM Tris-HCl pH 7.5, 10 mM MgCl₂, 2,5 mM DTT, and protease inhibitors) and filtered through Miracloth to remove debris. Crude extracts were centrifuged at 16,100 g to separate cytosolic, chloroplast, and mitochondrial fractions. The nuclear pellet, initially green due to residual chloroplasts, was washed repeatedly with LVM2 buffer (0.25 M sucrose, 1% Triton X-100, 10 mM Tris-HCl pH 7.5, 10 mM MgCl₂, 2,5 mM DTT, protease inhibitors, filtered through 0.25 μm pore size) until a white pellet was obtained, indicating removal of contaminating plastids. Purified nuclei were further enriched by sucrose cushion centrifugation in LVM3 buffer (2.5 M sucrose, 0.15% Triton X-100, 10 mM Tris-HCl pH 7.5, 2.5 mM MgCl₂, 2.5 mM DTT) and finally lysed in LVM lysis buffer (50 mM Tris-HCl pH 7.5, 10 mM EDTA pH 8, 1% SDS, protease inhibitors).

### sRNA-seq and bioinformatic analysis

Small RNA libraries with unique molecular identifiers (UMIs) were prepared and sequenced by Fasteris (Geneva, Switzerland), generating size-selected reads of 15-43 nucleotides. Raw data quality was assessed with FastQC (v0.11.9) and MultiQC (v1.12), and PCR duplicates were removed based on UMI sequences. Deduplicated reads were aligned to the *A. thaliana* reference genome (TAIR10, including mitochondrial and chloroplast sequences) and the ORMV genome using Bowtie (v1.3.1), allowing up to one mismatch and reporting up to 50 best alignments. BAM files were created with Samtools (v1.18) and sorted, indexed, and merged by size class (21-22 nt and 23-24 nt). Small RNA clusters were identified *de novo* using ShortStack (v3.8.5) (Axtell, 2013) and cluster-specific read counts were then quantified. Normalization factors were calculated from small RNA reads mapping to reference housekeeping RNAs (tRNAs, snoRNAs, rRNAs) and differential expression analyses were performed with DESeq2 (v1.46.0) (Love *et al*, 2014) in R (v4.2.1) using precomputed size factors. Log_2_ fold changes were shrunk with the R package ashr (Adaptive Shrinkage), and clusters with adjusted p-values < 0.05 and absolute log_2_ fold changes > log_2_ = 1.5 were considered significant. Data visualization and downstream analyses were performed with ggplot2 (Wickham, 2016) and GraphPad Prism10.

## Results

### Induced tolerance depends on CLASSY proteins

To investigate the mechanism through which Pol IV contributes to induction of tolerance, we studied plants deficient for CLASSY (CLSY) proteins, which are SNF2-domain proteins that recruit Pol IV to TEs leading to their silencing. Arabidopsis encodes four CLSY proteins (CLSY1/2/3/4) that each promote siRNA production at distinct genomic loci (Felgines *et al*, 2024; Yang *et al*, 2018; Zhou *et al*, 2022; Zhou *et al*., 2018). ORMV infection in *clsy1 clsy2 clsy3 clsy4* quadruple T-DNA (*clsy1234-TD*^Col^) or quadruple CRISPR/Cas9 (*clsy1234*-*C9*^Col^) knockout mutants (**Fig. S1**) leads to disease phenotypes that mimic those of *pol IV (nrpd1-3, pol IV-TD^Col^)* loss-of-function (**Fig. 2A**), indicating that CLSY-mediated recruitment of Pol IV to target loci is required for induced tolerance. Consistent with the upstream role of CLSY proteins in RdDM, both *clsy1234-TD*^Col^ and *clsy1234-C9*^Col^ quadruple mutants show loss of TE-derived 24 nt siRNAs, like *pol IV* mutants (**Fig. S2A**). Despite strong disease phenotypes after infection, the *pol IV* mutant and both *clsy1234* quadruple mutants accumulate virus levels similar to the wild type, showing that CLSY-Pol IV machinery is required for induction of tolerance (**Fig. 2B**).

**Fig. 2.**
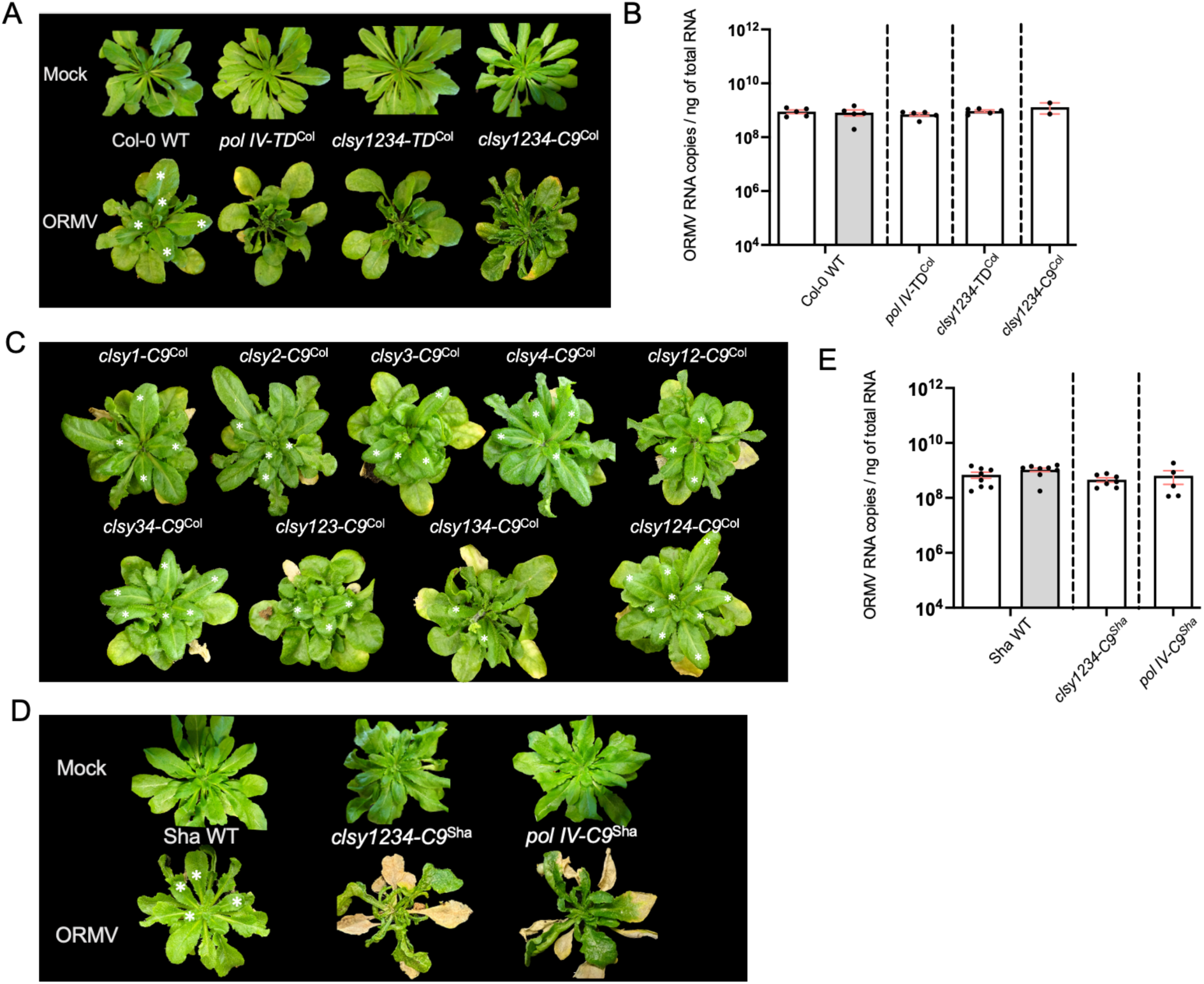
Disease recovery of ORMV-infected plants depends on CLASSY and Pol IV, and is independent of virus accumulation. (**A**) Phenotypes of mock-treated and ORMV-treated Arabidopsis Col-0 wild-type plants and *pol IV (nrpd1-3; pol IV-TD*^Col^) and *clsy1234* (*clsy1234-TD*^Col^ and *clsy1234-C9*^Col^) quadruple mutants at 28 dpi. (**B**) ORMV accumulation in apical leaves (RT-qPCR) of Col-0 and *pol IV* and *clsy1234* quadruple mutants at 28 dpi. (**C**) ORMV-infected single, double, and triple *clsy* mutants show normal disease recovery (asterisks). Triple mutants that include *clsy3* (*clsy123*, *clsy134*), exhibit mild disease recovery (less asterisks) whereas triple mutants only retaining CLSY3 (*clsy124*) exhibit enhanced disease recovery (more asterisks). Representative images were taken at 28 dpi. (**D**) Mock and ORMV-infected ecotype Sha WT and *clsy1234-C9*^Sha^ and *pol IV-C9^Sha^* mutants at 28 dpi. Unlike Sha WT plants, *clsy1234-C9*^Sha^ and *pol IV-C9^Sha^* mutants fail to develop disease recovery. (**E**) Accumulation of ORMV RNA in Sha WT and *clsy1234-C9*^Sha^ and *pol IV-C9^Sha^* mutants at 28 dpi. Viral RNA levels in (**B**) and (**E**) were determined by absolute quantification using TaqMan RT-qPCR. White columns, diseased leaves; grey columns, recovered leaves. Points represent individual biological replicates. Bars show the mean with standard error (SE).

Using a panel of Arabidopsis single, double and triple CRISPR/Cas9 mutants that knock out distinct combinations of *CLSY* genes, we demonstrated that CLSY3 plays a major role in the induction of tolerance and disease recovery (**Fig. 2C**). While mutations in individual *CLSY* genes (*clsy1, clsy2, clsy3 and clsy4*) or two *CLSY* genes (*clsy12, clsy34*) had no effect on recovery, mutations in three *CLSY* genes, including *CLSY3* (*clsy123* and *clsy134*), resulted in reduced disease recovery evidenced by fewer recovered leaves in the plant apex. Conversely, *clsy124* mutants that retain CLSY3 activity exhibited enhanced disease recovery, as shown by an increased number of recovered leaves, suggesting that the CLSY3 protein alone is sufficient for disease recovery in ORMV-infected plants. Analysis of Arabidopsis lines expressing β-glucuronidase (GUS) under control of the respective *CLSY* gene promoters (Zhou *et al*., 2022), showed that ORMV infection triggers strong CLSY3 activation (**Fig. S3**). Disease recovery is also observed in ORMV-infected Shadara (Sha) ecotype plants. Quadruple mutation of all four *CLSY* genes in the Sha ecotype (*clsy1234-C9*^Sha^) (**Fig. S4**) disrupts disease recovery like in the Col-0 ecotype (*clsy1234-C9*^Col^) (**Fig. 2A and Fig. 2D**), indicating that CLSY function in induced tolerance is not specific to the Col-0 genetic background. As shown in the *clsy1234-TD*^Col^ and *clsy1234-C9*^Col^ mutants, the *clsy1234-C9*^Sha^ mutant is also deficient for production of TE-derived 24 nt siRNAs (**Fig. S2, B**). Moreover, like for *pol IV* and *clsy* mutations in Col-0, the *pol IV* (*pol IV-C9^Sha^;* **Fig. S5**) and *clsy* mutations in Sha also have no effect on virus accumulation (**Fig. 2E**), confirming again that Pol IV and CLSY enable disease recovery by inducing tolerance rather than resistance.

### Induced tolerance is associated with siRNA synthesis from coding genes and transposons

To further determine the exact roles of DCL3, Pol IV and CLSY proteins in plant tolerance, we investigated the small RNA profiles in non-infected (mock) plants, and in diseased and recovered leaves of ORMV-infected plants at 28 dpi. The small RNA profiles of wild-type Col-0 plants were compared to those of *dcl3* (*dcl3-1*), *pol IV* and *clsy1234* (*clsy1234-TD^Col^)* null mutants. The virus-infected wild-type and mutant plants used for profiling showed the expected phenotypes (**Fig. S6, A**) and expressed the VSR protein (**Fig. S6, B**). Moreover, as observed above (**Fig. 2**), virus levels were similar in wild-type Col-0 and the mutants (**Fig. S6, C**). The profiling analysis revealed that wild-type plants accumulated abundant 23-24 nt siRNAs, as expected, whereas the small RNA profiles from *dcl3, pol IV* and *clsy1234* mutants contain abundant 21 nt siRNAs but relatively few 23-24 nt siRNAs (**Fig. 3A**). This data confirms expected molecular deficiencies in our mutants, which disrupt 23-24 nt siRNA biogenesis. Importantly, since *dcl3* mutants show ORMV disease recovery, this finding demonstrates that 24 nt siRNAs are dispensable for induction of tolerance. Instead, infection caused enrichment of endogenous 21 nt siRNAs in wild-type plants and all the tested mutants, most conspicuously in the *dcl3* line (**Fig. 3A**). The observation that *dcl3* plants exhibit enhanced disease recovery thus suggests a pivotal role of 21 nt rather than 24 nt siRNAs in induced tolerance.

**Fig. 3.**
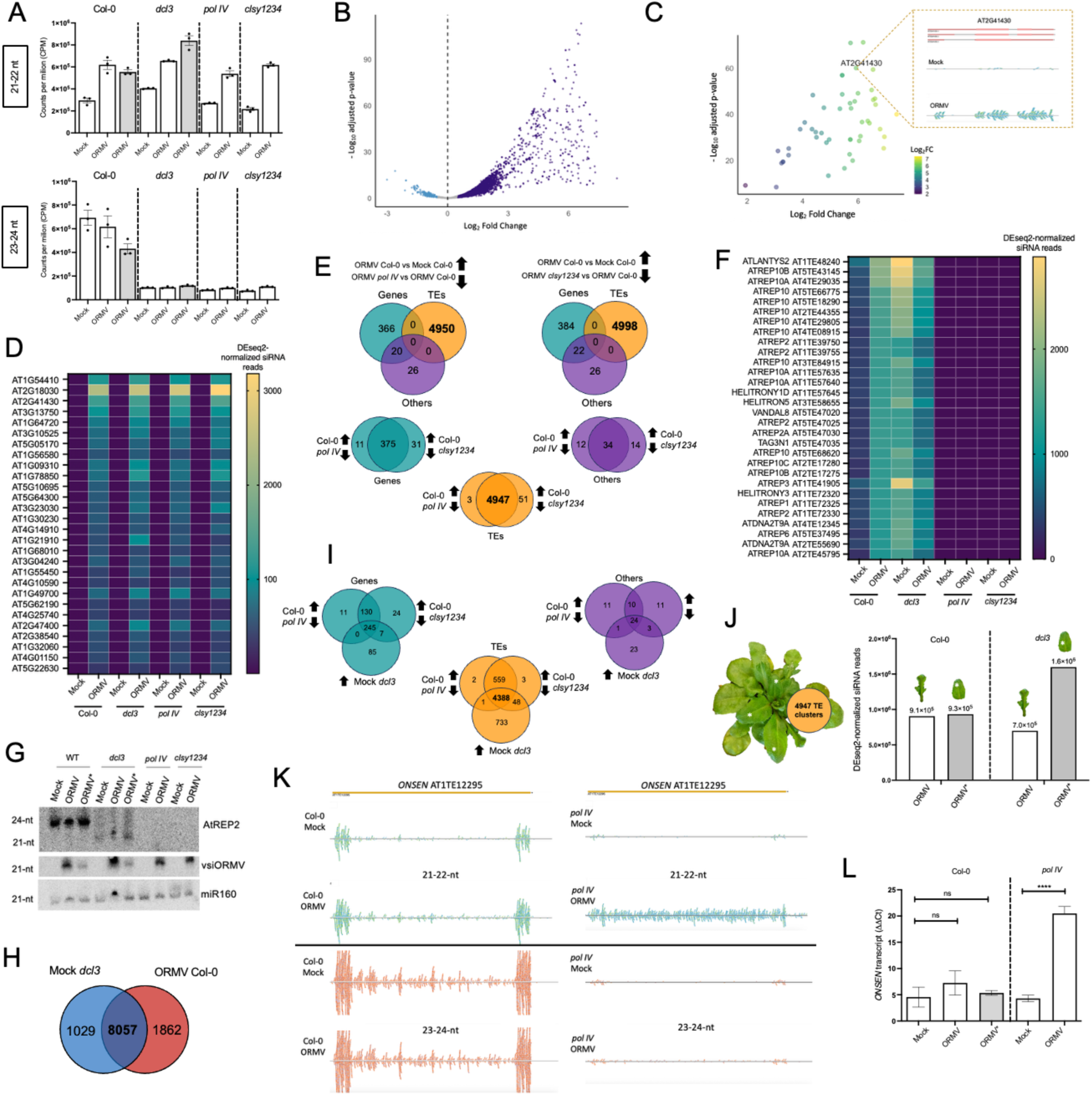
Virus-induced small RNA production from coding genes and TEs at 28 dpi. (**A**) Abundance of 21-22 nt and 23-24 nt small RNA reads in mock- or ORMV-treated Col-0 wild-type (WT) and mutant (*pol IV, clsy1234,* and *dcl3*) plants. **(B)** Activation of small RNA clusters upon ORMV infection. (**C**) Activation of vasiRNA clusters upon virus infection. (**D**) Virus-induced accumulation of vasiRNAs in Col-0 wild-type (WT) plants and mutants (*pol IV, clsy1234,* and *dcl3*). (**E**) Virus-induced siRNA clusters that are dependent on Pol IV and CLSY. The strong majority is associated with TEs (virus-associated TE-derived siRNAs, vaTE-siRNAs). (**F**) vaTE-siRNA accumulation in mock or ORMV-treated Col-0 wild-type plants and mutants (*pol IV, clsy1234,* and *dcl3*). (**G**) siRNA gel blot analysis confirming the accumulation of virus-derived siRNAs in infected tissues and the presence of 24 nt TE (AtREP2)-derived siRNAs in Col-0 WT plants. AtREP2-derived siRNAs are absent in *pol IV* and *clsy1234* mutants, whereas the *dcl3* mutant accumulates AtREP2-derived siRNAs that are 21 nt in length. (**H**) 21 nt siRNA clusters that are upregulated during viral infection in WT plants and already upregulated in *dcl3* plants prior to infection. (**I**) Pol IV/CLSY-dependent clusters upregulated in mock-treated *dcl3* plants. (**J**) Normalized reads for Pol IV/CLSY-dependent vaTE-siRNA derived from the same siRNA clusters identified in symptomatic and recovered tissues of Col-0 wild type (Col-0) plants and *dcl3* mutants. (**K**) siRNA mapping profiles across the *ONSEN* retrotransposon in mock and ORMV-treated Col-0 wild type (WT) and *pol IV* mutant plants. (**L**) Relative *ONSEN* transcript levels measured in Col-0 wild-type (Col-0) plants and *pol IV* mutants. White columns, symptomatic tissue; grey column, recovered tissue. Bars show the mean with standard error (SE).

Analysis of host-derived 21-22 nt siRNAs using ShortStack (Axtell, 2013) and DESeq2 (Love *et al*., 2014) revealed that ORMV infection caused a significant increase in the number of siRNA clusters mapped to the Arabidopsis genome (12,931 clusters with an adjusted p-value (padj) < 0.05; **Fig. 3B** and **Table S1**). Several of these clusters (748) mapped to coding genes and their strong induction upon infection (**Fig. 3C**) and other features fit to the definition of virus-activated siRNAs (vasiRNAs) (Cao *et al*, 2014). Typically, these vasiRNAs were 21-22 nt in length and mapped to exons of coding genes in both forward and reverse orientations (**Fig. 3C**), indicating that they originated from messenger RNAs (mRNAs) that were converted to double-stranded RNA (dsRNA) prior to DCL processing. vasiRNA synthesis in Arabidopsis was shown to require the SA-inducible RDR1 protein and DCL4 (Cao *et al*., 2014). Our small RNA profiles of ORMV-infected plants with different genetic backgrounds demonstrate that vasiRNA synthesis is independent of CLSY proteins, Pol IV and DCL3 (**Fig. 3D, Fig. S7, and Table S2**). By contrast, induced tolerance leading to disease recovery depends on Pol IV and CLSY, is repressed by DCL3, and does not require RDR1 or SA synthesis pathways (Kørner *et al*., 2018). We thus conclude that vasiRNAs, as defined above, are not required or sufficient for the induction of tolerance.

In order to identify small RNA profile changes that are essential for the induction of tolerance, we focused on the fraction of virus-induced 21 nt siRNAs that are Pol IV and CLSY-dependent (**Fig. 3E**). Of the 12,931 virus-induced 21 nt siRNA clusters identified in the Col-0 wild type, 4,947 clusters originated from TEs and were both Pol IV- and CLSY-dependent (**Fig. 3E**, **Fig. 3F**, and **Table S3**). The number of virus-induced clusters mapping to TEs and the highly induced number of siRNAs derived from these clusters (**Fig. 3F**) demonstrate that ORMV infection significantly stimulates the production of 21 nt siRNAs from TEs (virus-activated TE-derived siRNAs, vaTE-siRNAs).

In order to better understand the increased tolerance observed in *dcl3* mutants, we compared the 21 nt siRNA profiles of infected and mock-treated *dcl3* mutant plants. In this genetic background, TE-derived Pol IV and CLSY-dependent siRNAs accumulate as 21 nt rather than 24 nt products. Moreover, this strongly increased accumulation of 21 nt siRNAs is also observed in mock-treated *dcl3* plants (**Fig. 3G and Fig. S8**), also reflected in our genome-wide profiling data. Comparison of the 21 nt siRNA profile induced by ORMV in Col-0 with those of mock *dcl3* mutants showed that 8,057 virus-induced clusters are already upregulated in *dcl3* mock plants (**Fig. 3H**). Of these upregulated clusters in the mock-treated *dcl3* plants, the large majority are Pol IV and CLSY-dependent clusters of 21 nt TE-derived siRNAs (**Fig. 3H** and **Fig. 3I**). This overlap suggests that Pol IV and CLSY-dependent 21 nt vaTE-siRNAs associated with induced tolerance in infected plants are produced through non-canonical Pol IV-RDR2-DCL4 steps but are suppressed by DCL3 activity in the absence of infection. The lack of this negative regulation in *dcl3* mutants thus renders plants more competent for induced tolerance. The observation that virus infection of wild-type Col-0 plants causes reduced accumulation of 24 nt siRNAs derived from specific TEs, whereas the amount of 21 nt siRNAs derived from these same TEs is increased (**Fig. 3G** and **Fig. S7**) therefore suggests that the ability of plants to control DCL3 during infection to allow the production of 21 nt vaTE-siRNAs is critical for tolerance. The Pol IV/CLSY-dependent 21 nt vaTE-siRNAs, which are derived from almost 5,000 TE loci, are particularly abundant in recovered leaves of infected plants. This is particularly true for the recovered leaves of *dcl3* mutants, which show enhanced recovery (**Fig. 3J**). Taken together, these data support the specific role of vaTEsiRNAs in the induction of tolerance.

Our small RNA analyses also revealed virus-inducible 21 nt siRNAs arising from TE targets that are directly silenced by Pol IV (**Table S4**). This includes *ONSEN* Ty1/Copia long terminal repeat (LTR) retrotransposons, with these 21 nt siRNAs likely produced by DCL4 processing of virus-induced *ONSEN* Pol II transcripts (**Fig. 3K** and **Fig. 3L**). Past studies found that heat-responsive transcription factors bind motifs in *ONSEN* LTRs, causing *ONSEN* to be activated by heat stress (Cavrak *et al*, 2014; Ito *et al*, 2011). *ONSEN* insertions can provide adaptive responses on short evolutionary timescales by rewiring the expression of regulatory genes like *FLC* (Raingeval *et al*, 2024). Our data indicate that *ONSEN* is also activated by viral infection, perhaps due to infection favoring the alternative Pol IV-RDR2-DCL4 pathway and reducing the capacity of Pol IV-RDR2-DCL3 to produce 24 nt siRNAs for *ONSEN* silencing via RdDM (Ito *et al*., 2011). Production of 21 nt siRNAs from *ONSEN* is consistent with studies showing that transcriptionally reactivated TEs are repressed by RDR6-dependent mechanisms. The 21/22 nt siRNAs generated in this process can re-initiate RdDM independently of Pol IV (RDR6-RdDM). Subsequently, these TEs are again transcribed by Pol IV to re-initiate Pol IV-RdDM (Cuerda-Gil & Slotkin, 2016). The 21 nt virus-induced *ONSEN* siRNAs that we observe may arise via such a noncanonical RdDM pathway that re-silences *ONSEN* after an initial disruption of RdDM by virus infection. As disease recovery in ORMV-infected plants depends on RDR6 (Kørner *et al*., 2018), the RDR6-dependent production of *ONSEN*-derived 21 nt siRNAs may contribute to the induction of tolerance, alongside other RDR6-dependent secondary siRNAs.

### Role of the VSR and AGO1 during induced tolerance

Induced tolerance in ORMV-infected Arabidopsis Col-0 plants is restricted to carbon-sink tissues and coincides with the loss of the ability of the VSR to suppress GFP transgene silencing (Kørner *et al*., 2018). As tobamoviral VSRs interfere with RNA silencing and AGO-RISC assembly via small RNA sequestration (Csorba *et al*, 2007; Kurihara *et al*, 2007; Vogler *et al*, 2007), the concurrent loss of disease symptoms and GFP silencing in carbon-sink tissues during induced tolerance may result from virus-induced siRNAs competing with miRNAs and GFP-derived siRNAs for VSR binding. By interfering with the sequestration of miRNAs by the VSR, the virus-induced siRNA profile may prevent virus-induced alterations to leaf development, leading to disease recovery (Elvira-Gonzalez *et al*, 2025; Kørner *et al*., 2018). This model is supported by the observation that the levels of AGO1 were markedly reduced in diseased leaves but increased in recovered leaves (**Fig. 4A**), thus indicating dynamic regulation of AGO1 during infection and recovery. The low AGO1 levels in symptomatic leaves could be caused by miRNA sequestration by the VSR and the subsequent proteolytic degradation of unloaded AGO1 (Derrien *et al*, 2012; Hacquard *et al*, 2022), a mechanism conserved in mammals and flies (Martinez & Gregory, 2013; Smibert *et al*, 2013). The high AGO1 protein levels detected in recovered tissues would be consistent with 21 nt vaTE-siRNAs and other secondary siRNAs binding all available VSR protein, thereby competing to free host miRNAs to bind and stabilize AGO1.

**Fig. 4.**
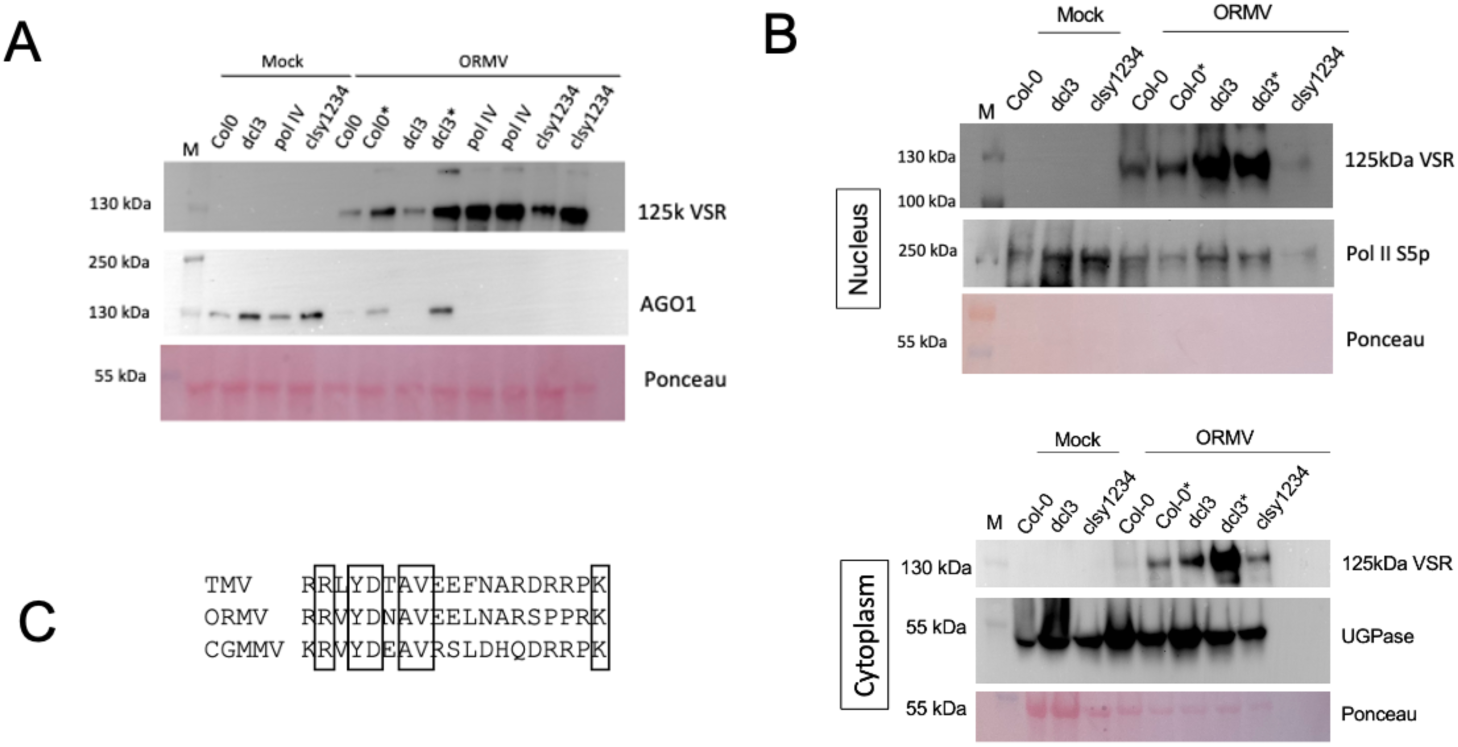
Protein accumulation in mock-treated or ORMV-infected plants. (**A**) Immunoblot analysis of AGO1 and the 125-kDa VSR of ORMV at 28 dpi. Ponceau staining for RuBisCo (51-58 kDa) is shown as a loading control. Samples from recovered tissues are indicated by an asterisk (*). (**B**) Nuclear and cytoplasmic accumulation of the 125 kDa VSR. Pol II S5p (B1 subunit of RNA polymerase II phosphorylated at Ser5) and UDP-glucose pyrophosphorylase (UGPase) are nuclear and cytoplasmic markers, respectively. Ponceau staining highlights cytoplasmic RuBisCo (51-58 kDa), which is absent in the nuclear fraction. (**C**) Sequence alignment of nuclear localization sequences (NLSs) in the VSR proteins of ORMV, TMV, and CGMMV.

Since AGO1-miRNA RISC assembly primarily takes place in the nucleus (Bologna *et al*, 2018), it is reasonable to ask how the VSR can compete with the nuclear AGO1 loading of miRNAs. Using cellular fractionation, we found that the ORMV VSR localizes to both the nucleus and cytoplasm (**Fig. 4B**). In fact, this ORMV VSR has residues homologous to the nuclear localization sequence (NLS) previously identified in TMV and cucumber green mottle mosaic virus (CGNNV) (dos Reis Figueira *et al*, 2002) (**Fig. 4C**). The VSR may therefore enter the nucleus and sequester miRNAs in the nucleus before AGO loading, thus interfering with AGO function and leading to alteration in plant development seen as disease symptoms, whereas vaTE-siRNAs and other 21 nt siRNAs that accumulate in sink tissues may saturate the neo-translated VSR in the cytoplasm before it enters the nucleus, thus allowing nuclear AGO1 loading with miRNAs and disease recovery (**Fig. 5**). Symptoms and recovery may thus be explained by the loss and restoration of AGO1 function, emphasizing the importance of AGO1 in determining the outcome of infection.

**Fig. 5.**
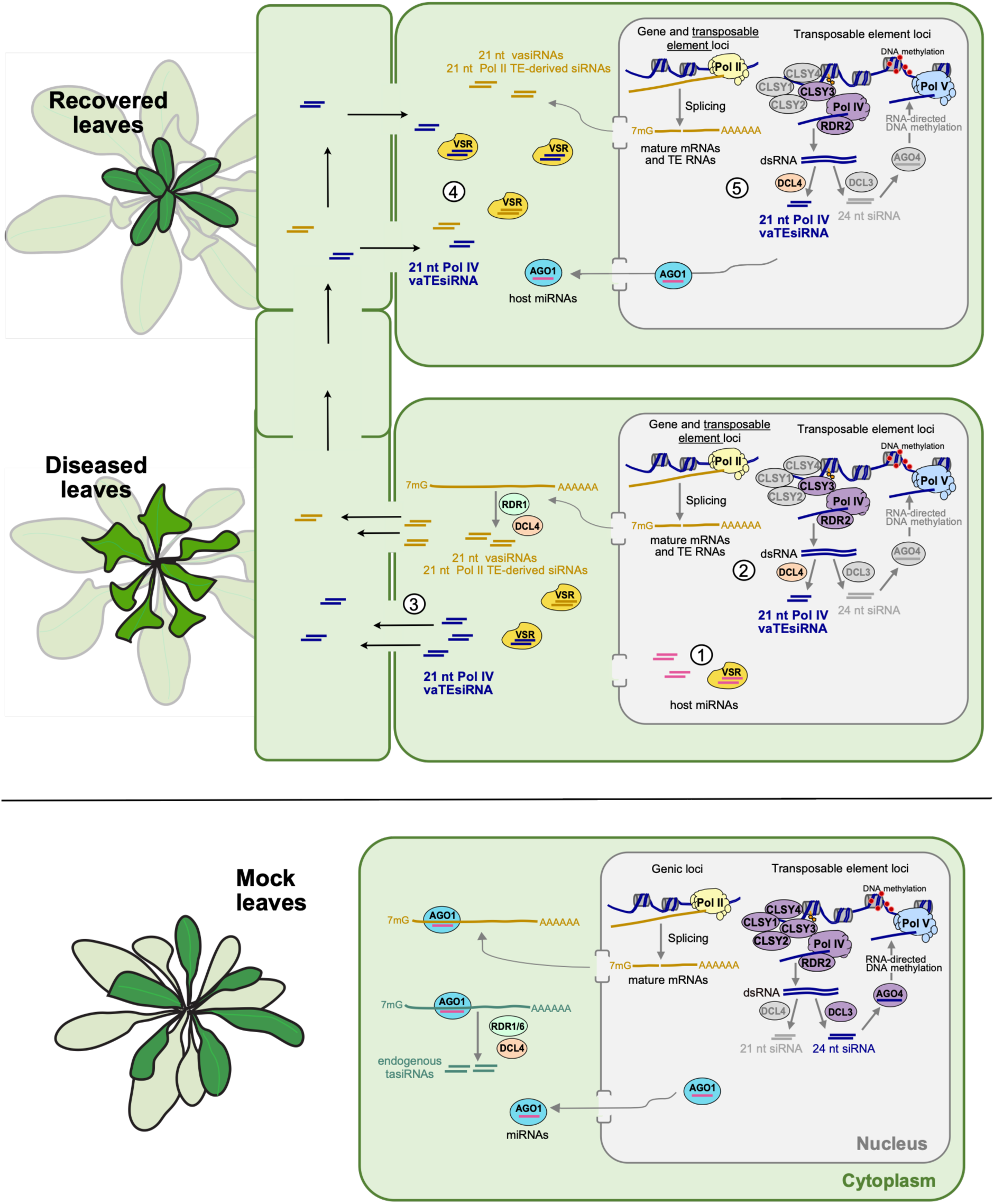
Small RNA and protein dynamics during disease and tolerance. Small RNA pathways in non-infected (mock) Arabidopsis rosettes: CLASSY (CLSY) proteins and RNA polymerase IV (Pol IV) mediate RNA-directed DNA methylation (RdDM) in the nucleus, whereas miRNA and other PTGS pathways act in the cytoplasm. **(1)** During ORMV infection the viral suppressor of RNA silencing (VSR) accumulates and moves to the nucleus where it binds miRNAs, interferes with miRNA loading of AGO1, and causes AGO1 degradation. **(2)** Infection leads to the induced production of siRNAs, particularly vaTE-siRNAs generated by a non-canonical CLSY-Pol IV-RDR2-DCL4 pathway, as well as of Pol II-dependent TE-derived siRNAs, and vasiRNAs. **(3)** Those siRNAs that are not captured by the VSR move systemically by source-sink flow to young sink leaves ahead of the virus. **(4)** In the young sink leaves, the accumulated siRNAs saturate the small RNA binding activity of the VSR in newly infected cells, thus interfering with the disease-inducing miRNA sequestration activity of the VSR. **(5)** During the course of infection in the newly infected leaf the virus again induces the production of vaTE-siRNAs and other classes of siRNAs, maintaining the accumulating VSR molecules in saturated state, thus maintaining tolerance.

## Discussion

Traditionally, plant-virus interactions have been viewed through the lens of resistance, emphasizing the activation of antiviral defenses that restrict viral replication and lead to low virus titers. However, resistance is inherently unstable in the face of the rapid evolutionary capacity of viruses, and thus cannot fully explain the widespread persistence of virus-host coexistence observed in nature. This coexistence is based on tolerance, a fundamental and mechanistically distinct defense strategy in plant-virus interactions (Pagan & Garcia-Arenal, 2020). Rather than inhibiting the virus, tolerance inhibits the negative impact of infection leading to disease. Tolerant plants are therefore capable of maintaining fitness while allowing high virus levels.

Using Arabidopsis plants infected with ORMV that undergo disease recovery via the induction of tolerance during development, we have demonstrated that the induction of tolerance is correlated with the virus-activated production of Pol IV-dependent 21 nt siRNAs encoded by TEs (vaTE-siRNAs). These TE loci may thereby play a pivotal role in mitigating the detrimental effects of viral infection on plant health. Production of the 21 nt vaTE-siRNAs is strongly enhanced in *dcl3* mutants, thereby explaining the enhanced tolerance induction in these mutant plants. The 24 nt siRNAs produced from Pol IV transcripts by DCL3 and the downstream components that use these siRNAs for RdDM are dispensable, as previously demonstrated (Kørner *et al*., 2018).

This study also demonstrates that upon induction of tolerance, DCL3 processing of Pol IV-dependent dsRNAs into 24 nt siRNAs is attenuated, increasing the levels of shorter 21 nt siRNAs. This is consistent with the previous notion that, in the absence of DCL3, Pol IV transcripts are processed by a nuclear DCL4 variant (Henderson *et al*, 2006; Panda *et al*, 2020; Pumplin *et al*, 2016). The occurrence of enhanced tolerance in *dcl3* mutants suggests that the Pol IV-RDR2-DCL4 pathway leading to tolerance is suppressed by DCL3 (**Fig. 5**). This places DCL3 at the core of nuclear regulation acting downstream of Pol IV and RDR2 to control the balance between viral infection and health. The potential inhibition of RdDM induced by displacement of DCL3 by DCL4 during processing of Pol IV-dependent transcripts in ORMV-infected plants may explain the observed activation of *ONSEN* retrotransposons. Such TE activation reveals an intriguing layer of virus-induced changes that could have wide-reaching impacts on the epigenome.

Mechanistically, we identified CLSY3 as a critical epigenetic regulator protein underlying induced tolerance. Of the four CLSY factors known to recruit Pol IV in Arabidopsis (Zhou *et al*., 2018), only CLSY3 was found to be transcriptionally induced by viral infection and sufficient to support disease recovery. These findings reveal the existence of a virus-inducible CLSY3-Pol IV-RDR2-DCL4-dependent siRNA pathway. In our model, CLSY3 targets specific TEs during virus infection, thereby producing 21 nt vaTE-siRNAs that mediate tolerance to the infection. CLSY and Pol IV-dependent 21 nt siRNA profiles overlap after virus infection, consistent with the CLSYs targeting Pol IV to specific genomic loci in a spatiotemporally regulated manner, as observed in healthy reproductive tissues (Xu *et al*, 2025; Zhou *et al*., 2022). Our small RNA-seq analysis identified TEs that could contribute to induced tolerance in ORMV-infected plants. Future studies may reveal whether this subset of TEs are targeted for Pol IV transcription through CLSY3 recruitment to DNA motifs via RIM factors, as recently shown for plant reproductive 24 nt siRNAs (Xu *et al*., 2025). Taken together, our work suggests that such TE loci encode information for production of 21 nt siRNAs, which can act to induce plant tolerance to pathogenic viruses.

Intriguingly, the key protein guided by 21 nt siRNAs, AGO1, shows a dynamic behavior in ORMV-symptomatic versus recovered tissues. Together with the detection of the VSR in the nucleus, this supports a dynamic model in which severe symptoms initially arise from miRNA-VSR interactions and AGO1 destabilization. These effects are subsequently counteracted by a massive accumulation of 21 nt siRNAs, predominantly generated through nuclear production of vaTE-siRNAs, which act as a molecular buffer against VSR-miRNA interference, thereby restoring RNA silencing activity and AGO1 function (**Fig. 5**). Taken together, our work expands the functional repertoire of TE-derived siRNAs, revealing their role as endogenous regulators of plant fitness during virus infection. Pol IV-dependent TE-derived siRNAs are known to silence stress-activated retrotransposons (Ito *et al*., 2011), re-silence spontaneously activated TEs via DNA methylation (Baduel *et al*, 2025) and control key steps in reproductive development (Chow & Mosher, 2023). Our study suggests that such siRNA-generating loci could also provide a reprogrammable response to pathogens. We suggest a conceptual framework in which tolerance emerges from the repurposing of the Pol IV-dependent small RNA biogenesis and PTGS pathways, offering a durable alternative to resistance-based plant protection strategies.

## Acknowledgments

This work was supported by the Agence National de la Recherche (grant ANR-21-CE20-0020-01 to MH, TB, VSL) and the European Union (grant HORIZON-TMA-MSCA-PF-EF 101108121 to LEG). We would like to thank Julie Law (Salk Institute) for providing the promoter-GUS fusion lines. We are grateful to S. Koechler and M. Alioua (AEG core facility, IBMP) for Illumina MiSeq validation of CRISPR/Cas9 constructs and Sanger sequencing identification of *A. thaliana* genomic deletions. We also thank Esther Lechner and Alexander Berr (IBMP) for kindly providing the AGO1 and Pol II S5p antibody, respectively.

## Author contributions

Conceptualization: MH, TB, VSL; Methodology: LEG, CH, CM, TB, MH; Investigation: LEG, CH, CM, LF, VF, PMK, JH; Visualization: LEG, CH, CM, LF, TB, MH; Funding acquisition: MH, TB, VSL, LEG; Project administration: MH, TB, VSL, LEG; Supervision: MH, TB; Writing – original draft: LEG, MH; Writing – review & editing: LEG, MH, TB, VSL

## Data, code, and materials availability

All data are available in the main text or the supplementary materials. The small RNA-seq data for this study have been deposited in the European Nucleotide Archive (ENA) at EMBL-EBI under accession PRJEB107732. (https://www.ebi.ac.uk/ena/browser/view/PRJEB107732). Materials are available through the corresponding authors upon request.

---

**Fig. S1.**
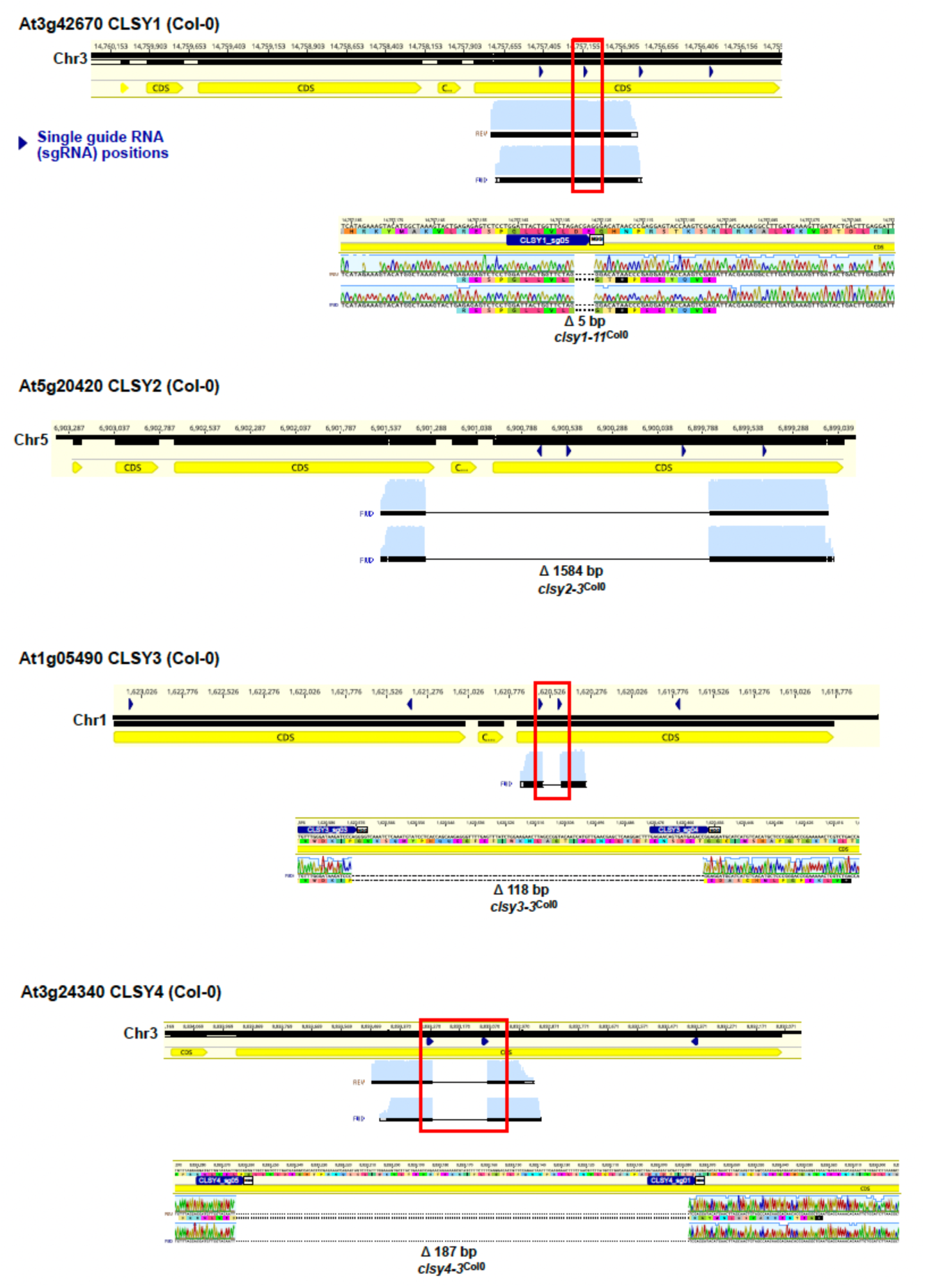
Generation and molecular characterization of *clsy* mutants in the *A. thaliana* Col-0 ecotype. Genomic organization of the CLSY1 (At3g42670), CLSY2 (At5g20420), CLSY3 (At1g05490), and CLSY4 (At3g24340) loci in *A. thaliana*. Coding sequences (CDS) are shown in yellow. Dark blue arrowheads indicate the positions of CRISPR/Cas9 single guide RNAs (sgRNAs) used for genome editing. Red boxes highlight the targeted regions of each gene and resulting deletions detected in the Sha ecotype. Sanger sequencing chromatograms shown below each locus confirm the mutations: a 5 bp deletion in *clsy1-11*^Col^, a 1584 bp deletion in *clsy2-3*^Col^, a 118 bp deletion in *clsy3-3*^Col^, and a 187 bp deletion in *clsy4-3*^Col^. Light blue profiles show the sequencing quality of bases aligned upstream and downstream of these genomic deletions.

**Fig S2.**
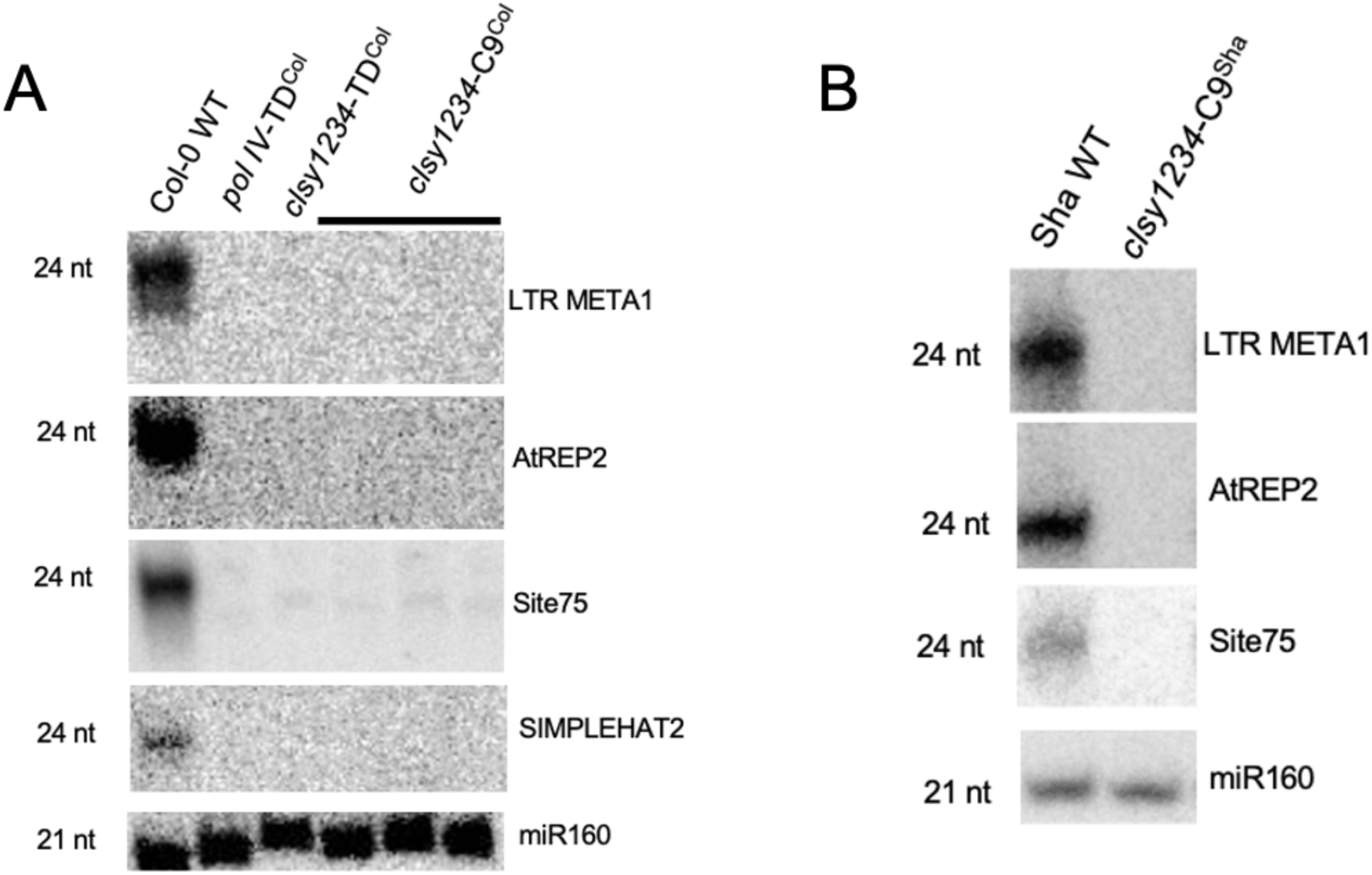
Accumulation of 24-nt siRNAs at selected loci in different Arabidopsis thaliana genotypes. **(A)** Northern blot analysis of 24 nt siRNAs derived from *LTR META1*, *AtREP2*, *Site75*, and *SIMPLEHAT2* in the Col-0 wild type and the indicated mutant backgrounds. **(B)** Northern blot analysis of 24-nt small RNAs derived from the same loci in Sha wild type and the indicated genotypes. Hybridization with a probe to detect miR160 was used as a loading control. The sizes of detected small RNAs (24 nt and 21 nt) are indicated at the left of each panel.

**Fig S3.**
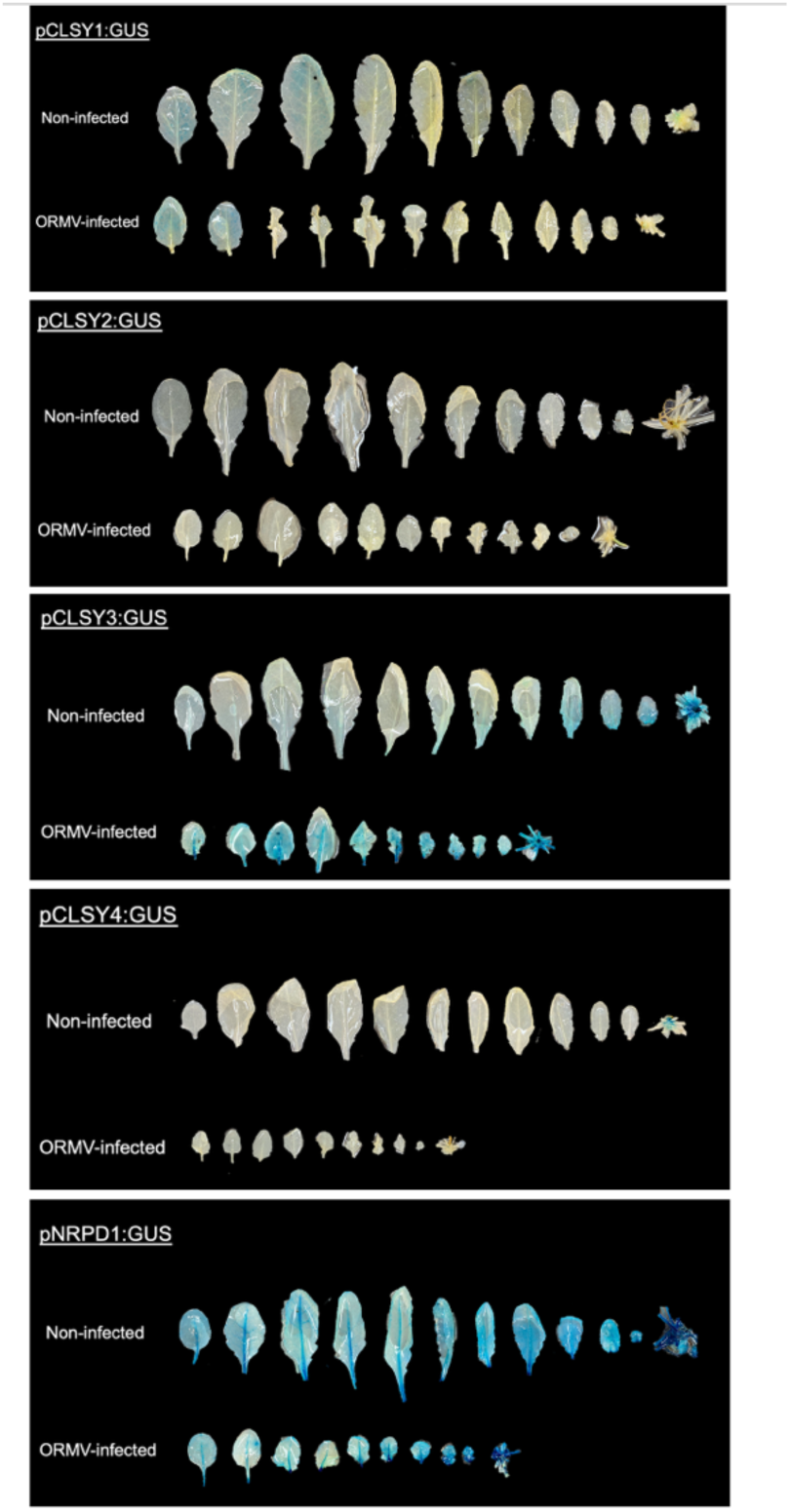
Histochemical analysis of *CLSY* gene-specific promoter:GUS fusion lines. p*CLSY1-*, p*CLSY2*, p*CLSY3*, p*CLSY4* or p*NRPD1*-driven GUS expression in mock and ORMV-treated plants at 21 dpi. Specific promoter activity is indicated by blue staining.

**Fig. S4.**
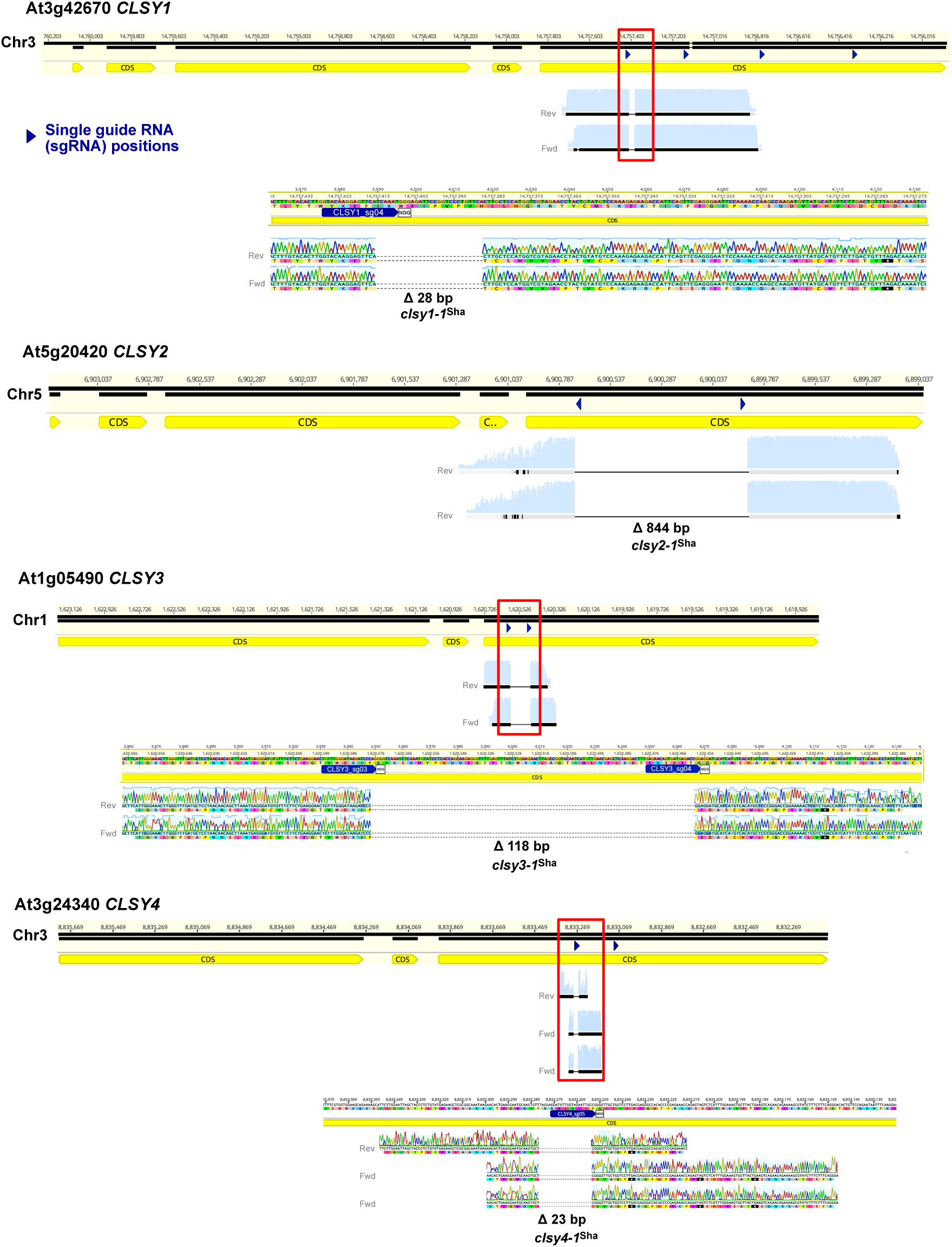
Generation and molecular characterization of clsy mutants in the *A. thaliana* Sha ecotype. Genomic organization of the CLSY1 (At3g42670), CLSY2 (At5g20420), CLSY3 (At1g05490), and CLSY4 (At3g24340) loci in *A. thaliana*. Coding sequences (CDS) are shown in yellow. Dark blue arrowheads indicate the positions of CRISPR/Cas9 single guide RNAs (sgRNAs) used for genome editing. Red boxes highlight the targeted regions of each gene and resulting deletions detected in the Sha ecotype. Sanger sequencing chromatograms shown below each locus confirm the mutations: a 28 bp deletion in *clsy1-1^Sha^*, an 844 bp deletion in *clsy2-1^Sha^*, a 118 bp deletion in *clsy3-1^Sha^*, and a 23 bp deletion in *clsy4-1 ^Sha^*. Light blue profiles show the sequencing quality of bases aligned upstream and downstream of these genomic deletions.

**Fig. S5.**
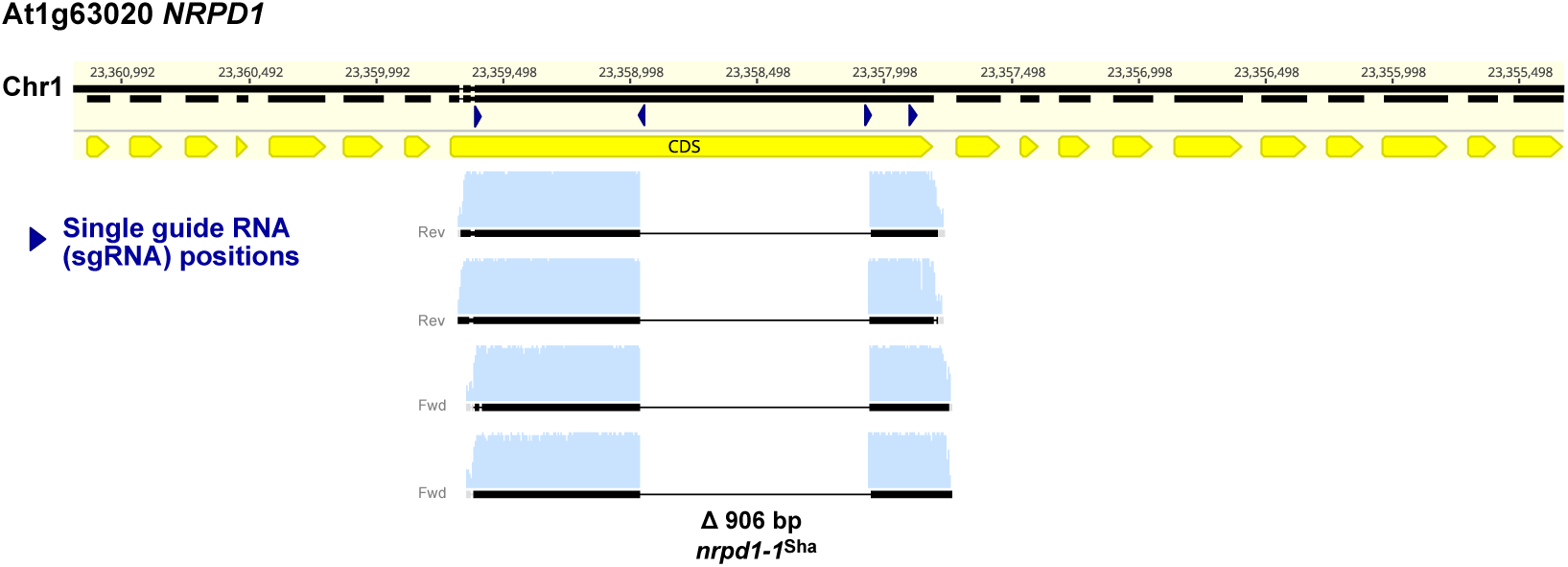
Generation and molecular characterization of a *nrpd1* mutant in the *A. thaliana* Sha ecotype. Genomic organization of the NRPD1 (At1g63020) locus in *A. thaliana*. Coding sequences (CDS) are shown in yellow. Dark blue arrowheads indicate the positions of CRISPR/Cas9 single guide RNAs (sgRNAs) used for genome editing. The targeted gene region and the 906 bp deletion mutation (*nrpd1-1^Sha^* / *pol IV-C9^Sha^*) detected in the Sha ecotype is shown. Sanger sequencing chromatograms below the *NRPD1* locus confirm this Cas9 mutation, which eliminates the entire active site region of the Pol IV enzyme. Light blue profiles show the sequencing quality of bases aligned upstream and downstream of these genomic deletions.

**Fig. S6.**
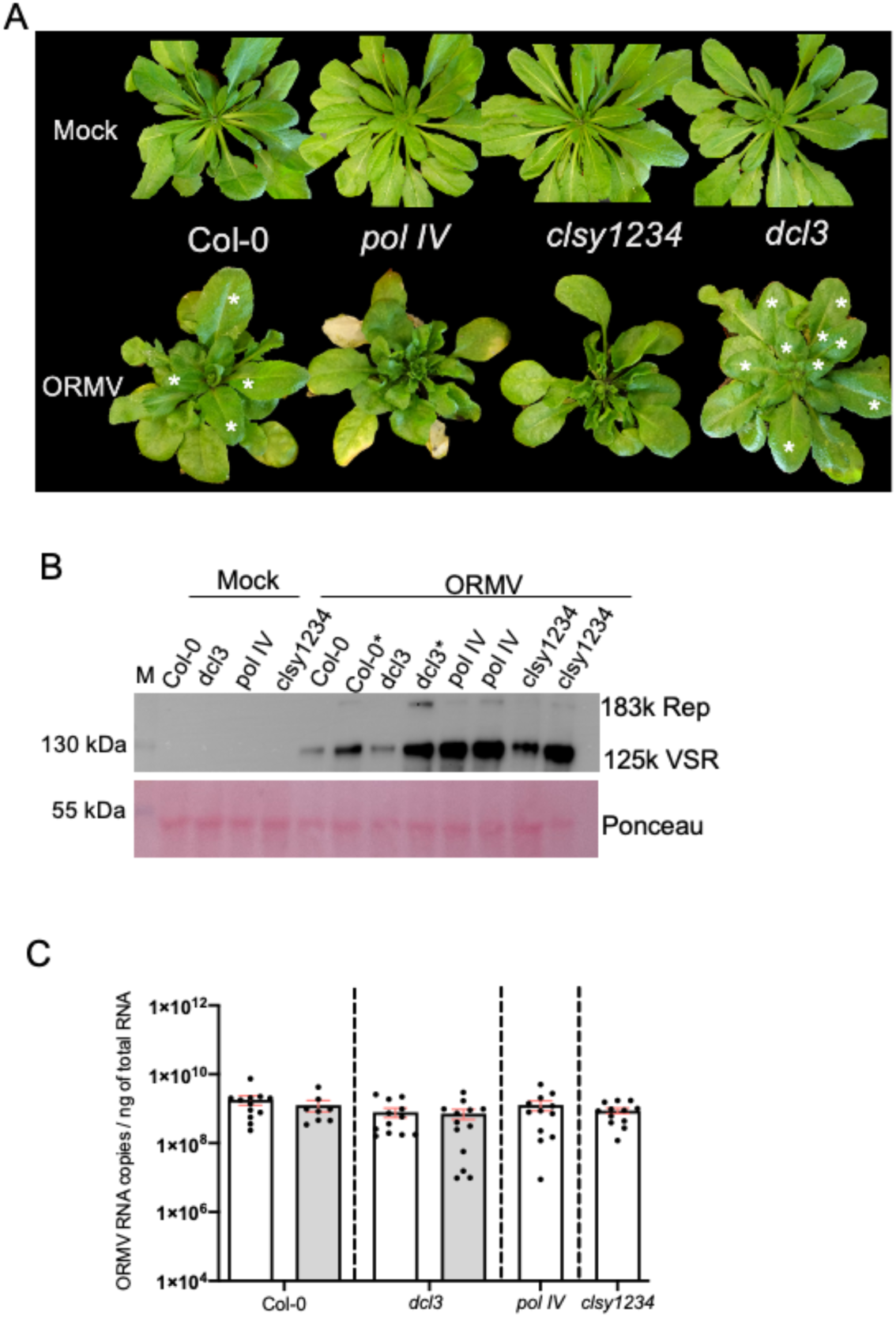
Samples for sRNAseq analysis. **(A)** Representative phenotypes of non-infected and ORMV-infected wild type plants (Col-O) and *pol IV*, *clsy1234*, and *dcl3* mutants at 28 dpi. Leaves showing disease recovery are indicated by asterisks. **(B)** Immunoblot detection of the ORMV 183 kDa replicase (Rep) and 125 kDa VSR proteins in apical leaves of plants showing disease (Col-0, *dcl3*, *pol IV*, and *clsy1234*) or disease recovery (Col-0 and *dcl3*) at 28 dpi. Samples from recovered leaves are labelled with an asterisk. Ponceau staining shows the large subunits of Ribulose-bisphosphate-carboxylase/oxygenase (RuBisCo; 51-58 kDa) as a marker for protein loading. M, protein size marker. **(C)** Virus accumulation in wild type plants (Col-0) and the mutants. Viral RNA levels were determined by TaqMan RT-qPCR at 28 dpi. Dots represent individual biological replicates. Mean values are shown with standard errors. Data from recovered leaf samples are shaded in grey.

**Fig. S7.**
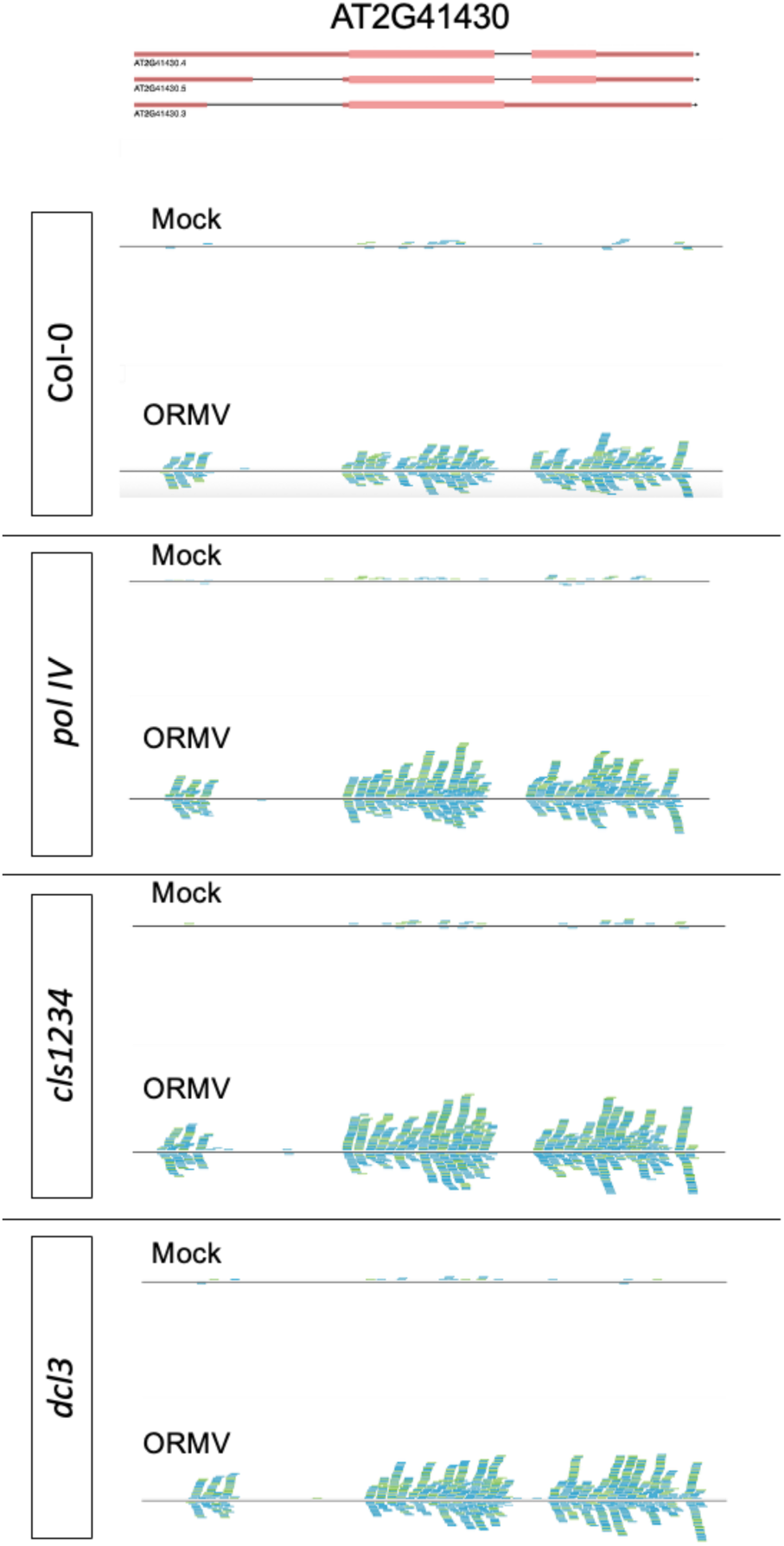
Virus-activated small interfering RNAs (vasiRNAs). vasiRNAs mapped to the gene of origin (AT2G41430) in the Col-0 wild type and mutants (*pol IV*, *clsy1234* and *dcl3*).

**Fig. S8.**
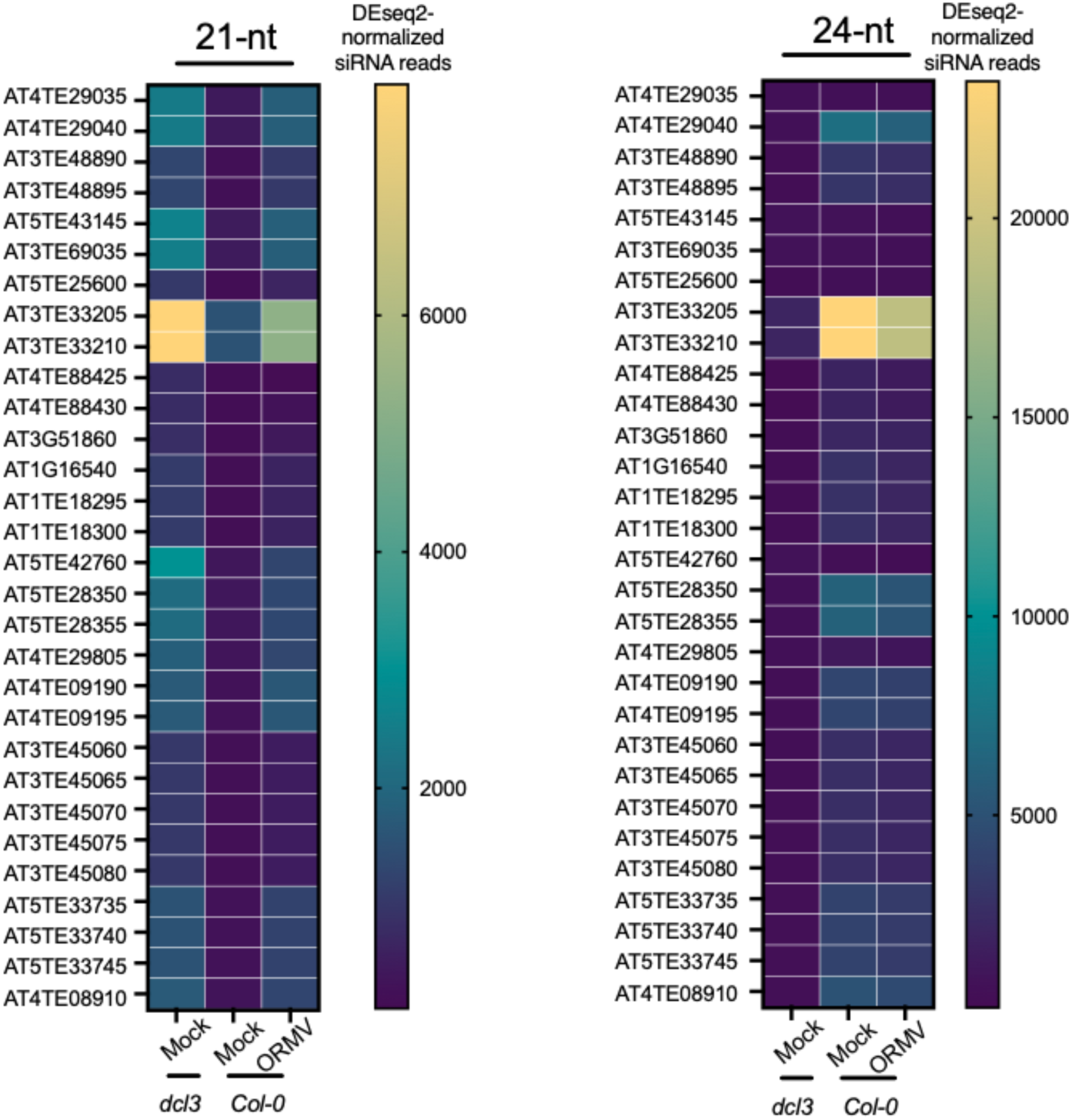
Regulation of TE-derived siRNAs. TE-derived siRNAs that are upregulated by virus infection in the Col-0 wild type are also upregulated in the non-infected, mock-treated *dcl3* mutant. Whereas 21 nt TE-derived siRNAs are upregulated upon infection, the 24 nt TE-derived siRNAs are down-regulated.

## Table legends

**Table S1.** Virus-induced siRNA clusters in Col-0 wild type. DEseq2-normalized siRNA reads (adjusted p-value (padj) < 0.05).

**Table S2.** Virus-induced vasiRNA clusters in Col-0 wild type and mutants (*pol IV, clsy1234* and *dcl3*). DEseq2-normalized siRNA reads of three biological replicates per condition (averaged reads). Log2FC > 1.9; adjusted p-value.

**Table S3.** Virus-induced, Pol IV and CLSY-dependent TE-derived siRNA clusters. DEseq2-normalized siRNA reads of three biological replicates per condition (averaged reads).

**Table S4.** Virus-induced TE-derived siRNA clusters that are independent of Pol IV and CLSY. DEseq2-normalized siRNA reads of three biological replicates per condition (average reads).

**Table S5.** List of primers and probes used in this study

